# Canonical BMP2/4 signaling degrades SOX2 in glioblastoma propagating cells

**DOI:** 10.1101/2022.02.03.479056

**Authors:** Flavio Cimadamore, Alejandro Amador-Arjona, Chen Farhy, Chun-Teng Huang, Harley Kornblum, Alexey V. Terskikh

## Abstract

The effect of Bone Morphogenetic Proteins (BMPs) on glioblastoma propagation cells (GPCs) is being debated [1-5]. We observed that exposure of GPCs to BMP2/4 initiates a rapid proteasomal degradation of the pluripotency factor SOX2 followed by the differentiation of GPCs into glial cells using 5 primary human glioblastoma lines tested. Enforced expression of SOX2 in GPCs antagonized BMP2/4-induced gliogenesis and reduction of proliferation. Both outcomes are consistent with the role of SOX2 in normal neurogenesis [6-8]. Using gene expression analysis, we observed that level of SOX2 in different glioblastoma lines tracks with the levels of oncogenes H/K/N-RAS and OLIG2. Our work provides support for BMP2/4-SOX2 axis in GPCs and points to the proteasome-regulated degradation of SOX2 that initiates glycogenic differentiation similar to that observed during normal development [6, 7]. Our work supports a potential for developing novel therapeutic strategies aimed at differentiation of GPC into non-tumorigenic glia.

## Introduction

Glioblastoma represents the most common type of intracranial tumor (50% of total gliomas) and, with an incidence of approximately 10000 new cases each year in the United States [9] and a very poor prognosis (median survival = 12-15 months) [10], it is undoubtedly one of the most dreaded types of brain tumors. The standard therapeutic approach for the treatment of this disease is radiotherapy in combination with chemotherapeutic Temozolomide treatment. Despites such treatments, glioblastoma are inexorably recurrent, underscoring the dire need for new, more efficient therapies. Glioblastoma is heterogeneous and a growing body of evidence speaks in favor of the existence, within the tumor mass, of a subpopulation of highly undifferentiated stem-like cells that resemble normal neural stem cells (NSCs) in terms of marker expression and differentiation potential [11-13]. Such population, termed Cancer Stem Cells or Tumor Initiating Cells, or Tumor Propagating Cells. Glioblastoma Propagating Cells (GPCs, the term we will use here), efficiently forms tumors in xenograft experiments and seem to be responsible for the radio- and chemo-resistance to standard clinical treatments which inexorably lead to GBM recurrence [14, 15]. Recent large-scale single-cell RNA-seq analyses of freshly isolated glioblastoma tumors revealed trilineage hierarchy centered on cancer progenitors, which appear to be the equivalent of glial progenitors during normal brain development [16]. Remarkably, glioblastoma progenitors were identified as the most proliferating cells within tumor mass driving chemoresistance and tumor growth. RNA velocity analysis revealed conservative cellular trajectories from GPCs to differentiated neuronal, oligo, astrocytic, and mesenchymal cells within all primary tumors analyzed [16]. Such hierarchical organization and naturally occurring differentiation trajectories in human glioblastoma identifies proliferating GPCs as target cells and strongly support the idea of further promoting naturally occurring differentiating of glioblastoma GPCs into non-tumorigenic cells.

Developmental pathways controlling self-renewal and differentiation of normal NSCs may be relevant to GPCs. For instance, the transcription factor SRY (sex-determining region)-box 2 (SOX2), which is required for self-renewal, proliferation and neurogenic differentiation of different type of NSCs [6, 7, 17, 18], is highly expressed *in vivo* in hyperproliferative areas of glioblastoma [19], marks proliferating GPCs ex vivo [16], expressed, albeit to variable extent, in cultured GPCs [5, 11] and functions as an oncogene by supporting their proliferation and tumor initiation capability [20]. On the other end, Bone Morphogenetic Proteins (BMPs), a group of cytokines of the TGF-β superfamily implicated in differentiation, apoptosis and proliferation of NSCs according to their developmental stage and microenvironment [21, 22], have been shown to exert a tumor suppressor activity on glioblastoma due to their ability to promote differentiation of GPCs and loss of their tumorigenic potential [1-3]. Despites the identification of such important players, the molecular mechanisms underlying SOX2 and BMP functions in GBM GPCs is poorly understood. Recent work from two independent groups arrived at different conclusions with respect to BMP2/4 effect on glioblastoma TPC. Sachdeva et al., 2019 suggested that exposure to BMP2/4 induces glioma stem cell quiescence [4]. Remarkably, pretreatment glioblastoma with BMP4 led to 3-4 times increase in survival upon orthotopic transplantation in mice [4] suggesting a dramatic improvement compared to the current clinical standards. In contrast, Dalmo et al., 2020 concluded that response is heterogenous among 40 primary human glioblastomas analyzed [5]. Invariably, such response in sensitive cells induced SMAD1/5/9 phosphorylation, SMAD4 expression, and consistent downregulation of SOX2 [5]. Our own observations align closely with the work of Dalmo et al., 2020. We observed that exposure of glioblastoma TPCs (5 primary lines) to BMP2/4 resulted in rapid downregulation of SOX2 followed by reduction in Ki67 and induction of GFAP (Fig. 3A). GBM3 line was initially unresponsive. We discovered that GBM3 was lacking expression of BMPR1; transduction with BMPR1restored the phenomena. We discovered that BMP2/4 initiates proteasomal degradation of SOX2. Enforced expression of SOX2 in glioblastoma TPCs rescued BMP2/4-induced gliogenesis and reduction of proliferation. It is reassuring that both outcomes are consistent with the role of SOX2 in normal neurogenesis previously uncovered by others and our group [6-8]. Further, SOX2 may be required in GSCs to maintain high levels of known oncogenes such as RAS proteins with NRAS being the oncogene more susceptible to loss of SOX2. These results provide support for BMP2/4-SOX2 axis in glioblastoma TPCs and points to the proteasome-regulated degradation of SOX2 that initiates glycogenic differentiation similar to that observed during normal development [6, 7].

## Results

### Downregulation of SOX2 is mandatorily required for BMP-induced differentiation of GPCs

Low passage cultures of human primary glioblastoma in NSC media (i.e. serum-free media in the presence of bFGF and EGF) robustly enriches for GPCs [11]. We exploited such observation to derive low passage GPC lines from several primary glioblastoma patients. As expected, our cultures expressed high levels of markers common to both NSCs and GPCs such as Nestin and SOX2 (Fig. S1A). We also verified their tumor initiation capability by transplanting luciferase-transduced GPCs into immunodeficient Rag2-/- & common γ-chain -/-knockout mice. Tumors could be observed after 43 days with as little as 1000 transplanted cells (Fig. S1B). To evaluate the efficacy of BMP treatment, we treated the cells with BMP2 or BMP4 for 3 days. Even in this short differentiation paradigm, BMP2/4 induced downregulation of the pluripotency factor SOX2, loss of proliferation potential (as evaluated by staining for the cycling marker Ki67) and acquisition of a glial fate (staining for the astrocyte marker GFAP) in 4 out of 5 patient-derived GPCs (Fig. 1A, Fig. S2). For comparison, growth factors withdrawal (No GFs conditions) didn’t induce appreciable differentiation (Fig. S2). Notably, TGFβ1, a closely related cytokine belonging to the same superfamily, didn’t induce SOX2 downregulation or loss of proliferation and only marginally increased GFAP expression in comparison to growth factor withdrawal (Fig. S2). even when used at concentrations 100-fold higher than its Effective Dosage 50 (ED50). Since neither growth factor withdrawal nor TGFβ1 could reproduce BMP2/4 effect on GPCs, the BMP2/4 effect is specific and likely mediated through BMPR complexes. We assessed the effect of SOX2 knockdown in GPCs and noticed that shRNA against SOX2 was sufficient to mimic BMP2/4 treatment, namely, inducing loss of Ki67 and increase of GFAP expression (Fig. 1B-E). This BMP2/4 differentiation effect was observed even when the cells were cultured in self-renewing conditions (i.e. in the presence of bFGF/EGF), optimized to prevent differentiation [11]. This points to the dominant effect of BMP2/4 signaling over bFGF/EGF signaling and suggests that reducing SOX2 levels is sufficient to mediate GPC differentiation. The ability of BMPs to downregulate SOX2 in such a short time and the fact that loss of SOX2 mimicked BMP treatment prompted us to investigate the link between these two important regulators of GPC biology.

**Figure 1:**
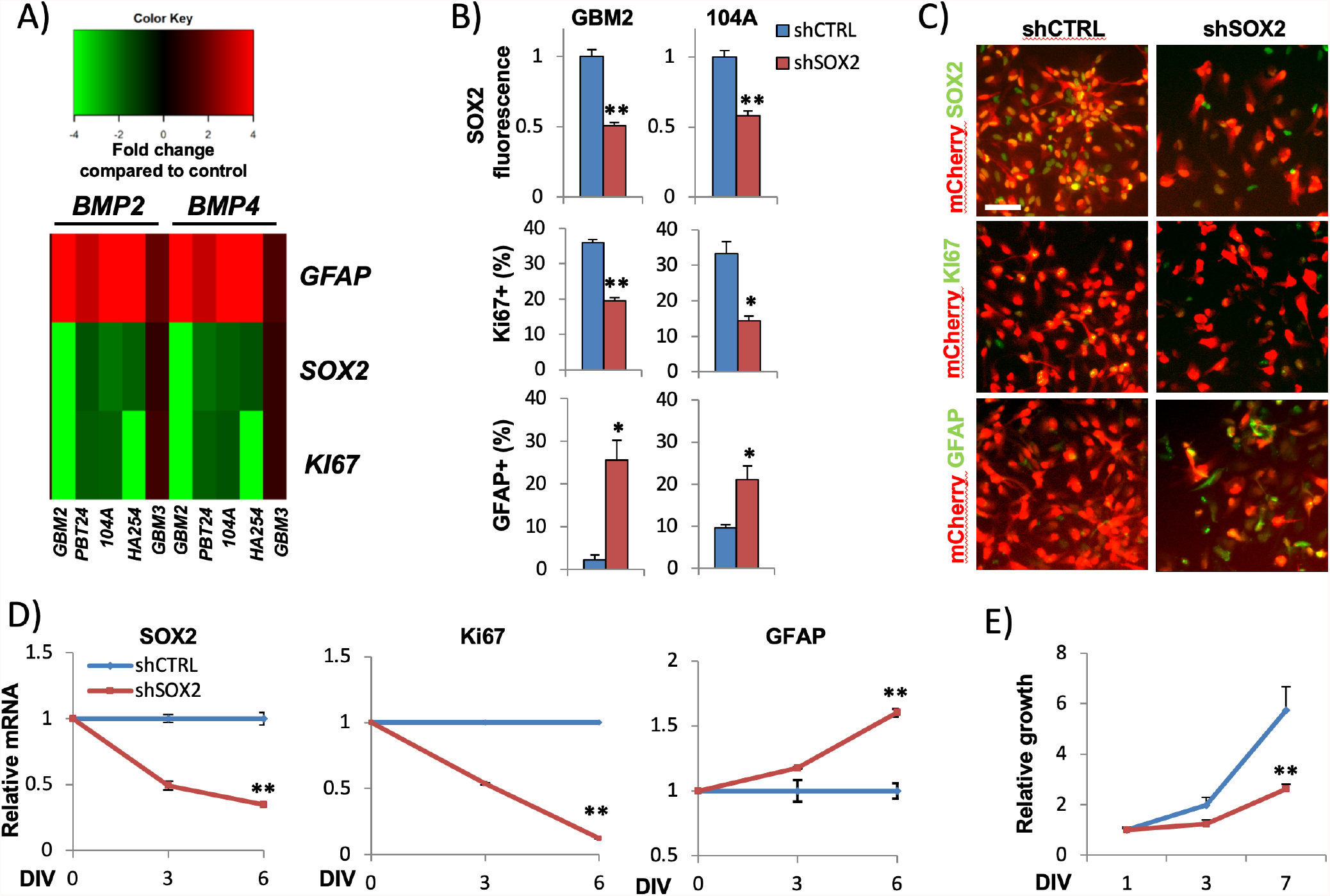
BMPs and SOX2 knockdown reduce proliferation and induce gliogenic differentiation of GPCs. A) Heatmap summary of BMP effect on several GPCs after 3 days of treatment. Values are fold change in marker expression (GFAP, KI67, and SOX2) in comparison to control (no growth factors) as assessed by immunocytochemical quantification. B-C) Quantification (B) and Immunocytochemistry (C) for SOX2, KI67 and GFAP in GBM2 and 104A GPC lines after 7-10 days of SOX2 knock-down induction. mCherry+ cells stably express either doxycycline-inducible scramble control shRNA (shCTRL) or SOX2-specific shRNA (shSOX2). D) Quantitative-PCR confirming SOX2 and KI67 downregulation and GFAP increase in inducible shSOX2-GBM2 GPCs 6 days after doxycyline treatment. E) Growth curve of shSOX2 GBM2 showing reduced proliferation in comparison to control cells. DIV=days in vitro; error bars = standard error; *p<0.05; **p<0.005. Scale bars = 50*µ*m

To understand if gliogenic differentiation and loss of GPC proliferation potential is direct consequence of SOX2 ablation downstream of BMP2/4, we overexpressed exogenous SOX2 under the control of a strong promoter (human phosphoglycerate kinase - PGK) and then treated GPCs with BMP2/4. Remarkably, enforces level of SOX2 antagonized BMP2/4 effect (Figure S3) preventing the loss of proliferation and astrocytic differentiation (Fig. 2A-B, Fig. S4A-B). These results suggest that the loss of SOX2 is a primary event downstream of BMP2/4 and it is required to mediate BMP2/4 effects on GPCs.

**Figure 2:**
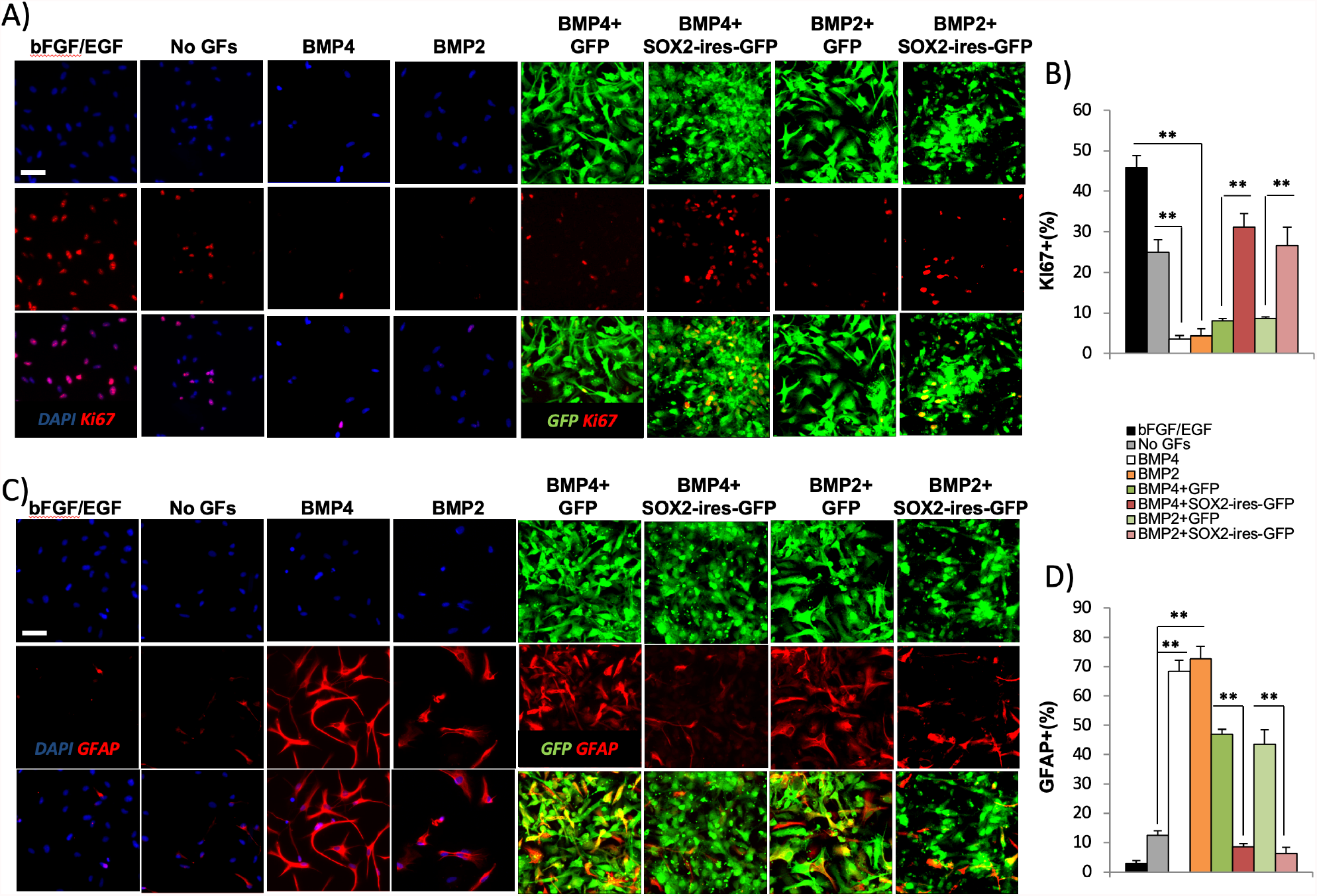
SOX2 overexpression blocks BMP-induced loss of proliferation and increased gliogenesis in GPCs. A-B) Immunostaining (A) and quantification (B) of KI67 in BMP2/4 treated GBM2 GPCs (3 days of treatment) infected with pgk-GFP or pgk-SOX2-IRES-GFP lentiviruses. Wild type cells treated with bFGF/EGF, no growth factors (NoGFs) or BMP4 are included for comparison. Note that SOX2 overexpression rescues the number of KI67+ cycling GPCs. C-D) Immunostaining (A) and quantification (B) of GFAP in BMP2/4 treated GBM2 GPCs (3 days of treatment) infected with pgk-GFP or pgk-SOX2-IRES-GFP lentiviruses. Note that SOX2 overexpression abolishes BMP-mediated increase in gliogenesis. Error bars = standard error; **p<0.005. Scale bar = 50*µ*m

### BMPR1 and SMAD phosphorylation is required for SOX2 degradation

Out of the five patient-derived GPC lines assayed for BMP activity, one (GBM3) didn’t downregulate SOX2 in response to BMP2/3 treatment (Figures 1A,4A). Similarly, GBM3 didn’t exhibit loss of proliferation (Fig. 1A), differentiation (Fig. 1A) and activation of BMP downstream effectors pSMAD1,5,8 (Fig. 3B-C). Of note, this line was still sensitive to SOX2 knockdown because a 40-50% downregulation of SOX2 using shRNA (Fig S6A, B, E) was sufficient to reduced proliferation (Fig. S6C-F). To understand this naturally occurring resistance to BMP2/4, we profiled the expression of known BMP receptors by qPCR. In contrast to other BMP-responsive lines, GBM3 clearly lacked the expression of BMPR1A (Fig. 3D). We introduced BMPR1A into GBM3 by electroporation and assessed responsiveness to BMP4 b. Remarkably, the exogenous expression of BMPR1A was sufficient to re-establish a BMP2/4 sensitive and downregulation of SOX2 (figure 3E-F). These results strongly suggest that BMPR1A mediates the observed SOX2 downregulation at least in GBM3 line.

**Figure 3:**
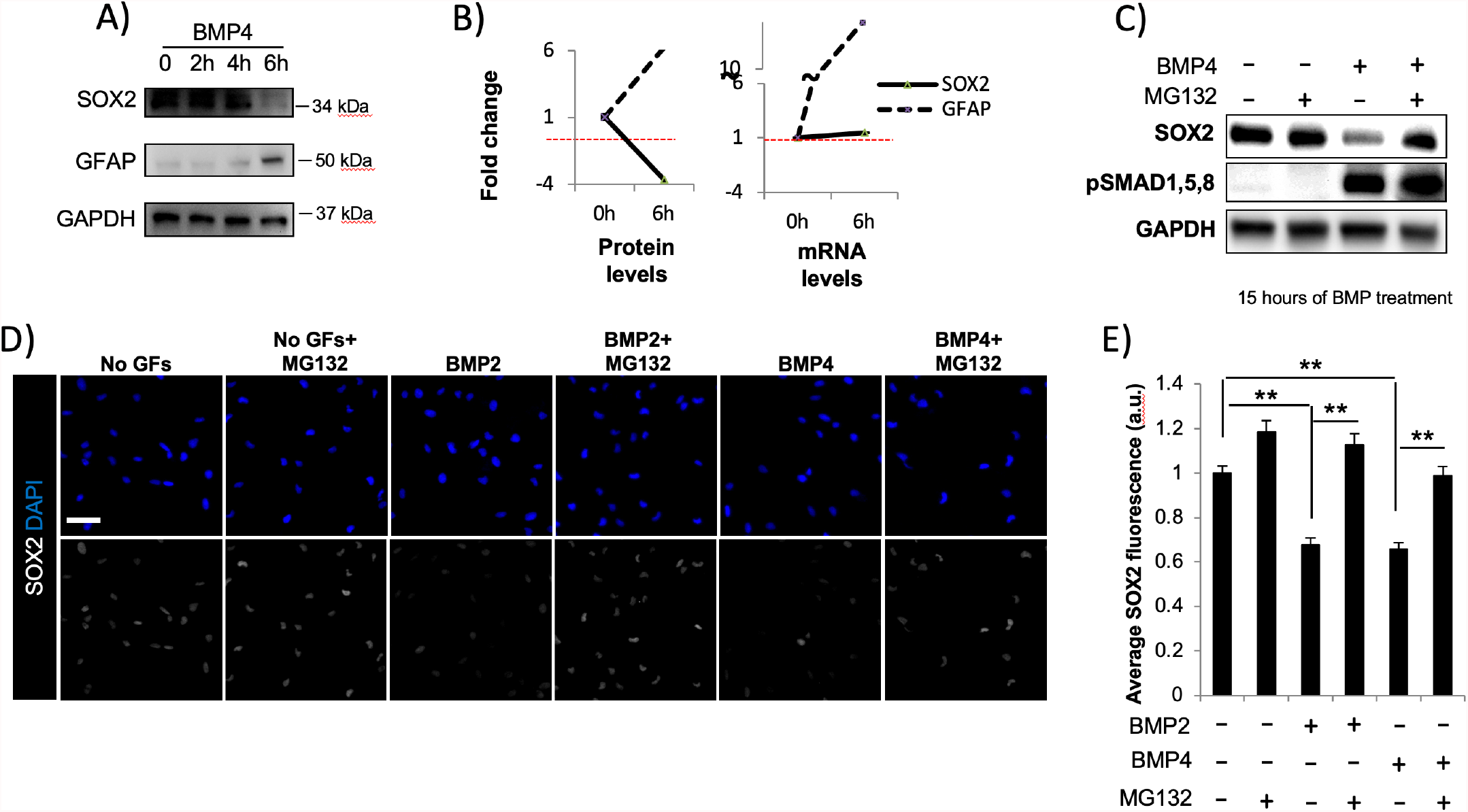
BMP-mediated degradation of SOX2 requires a functional proteasome system. A) Western blot showing the kinetics of SOX2 and GFAP expression during the first 6 hours of BMP4 treatment in GBM2 GPCs. B) Quantification of SOX2 and GFAP protein (left graph) and mRNA (right) levels before (0h) and after 6 hours (6h) BMP4 treatment. Note that in contrast to the protein, SOX2 mRNA at this early time point is not yet downregulated. C) Western blot for SOX2 and pSMAD1,5,8 in GBM2 GPCs cultured under the reported conditions for 15h. Note that addition of MG132 maintain high levels of SOX2 in the presence of BMP4. D) Immunostaining for SOX2 in GBM2 GPCs cultured under the reported conditions for 18 hours. Again, note that MG132 rescues BMP-mediated SOX2 degradation. E) Quantification of average SOX2 levels shown in (D). Error bars = standard error; **p<0.005. Scale bars = 50*µ*m

**Figure 4:**
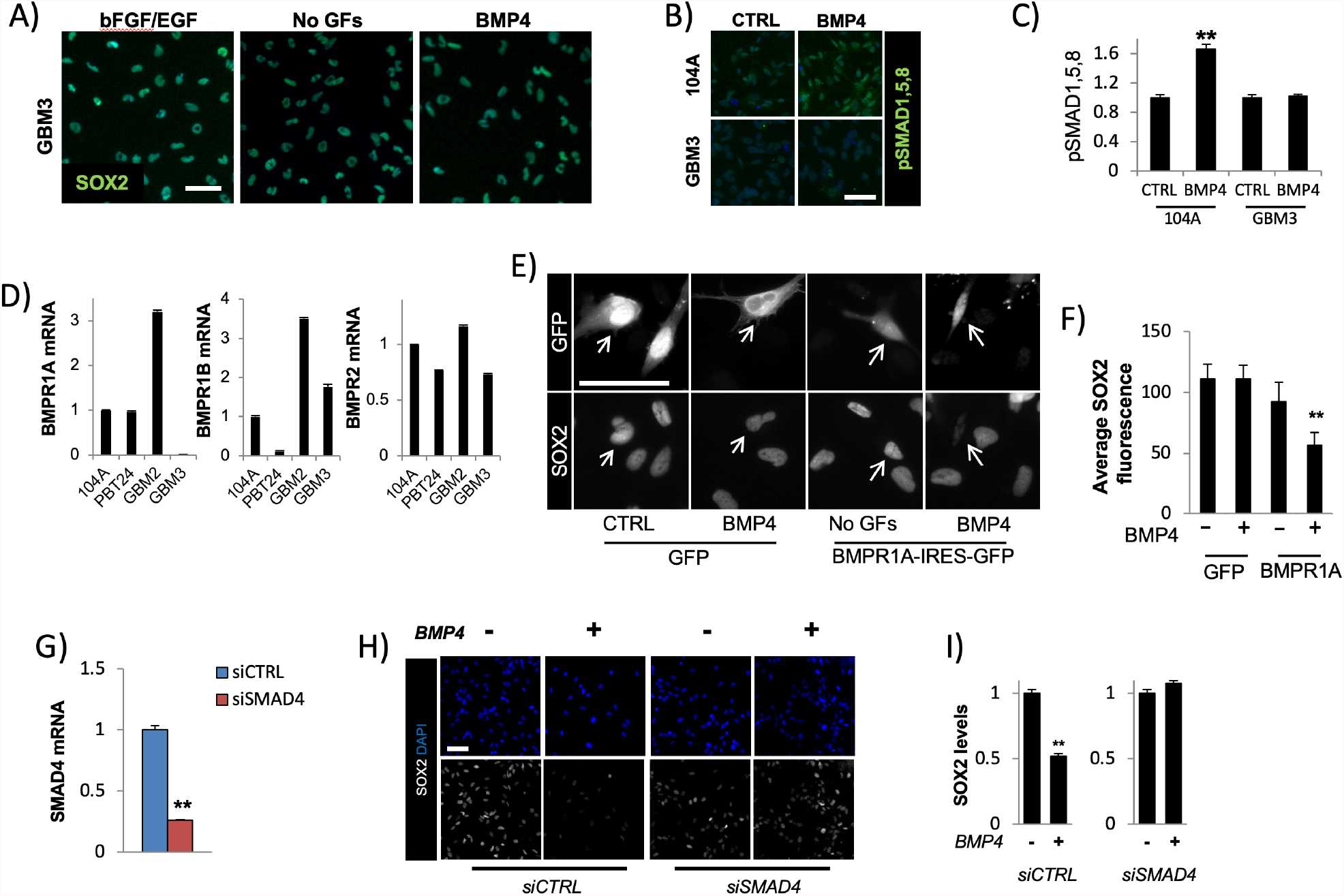
A canonical BMP signaling mediates SOX2 degradation. A) Immunostaining for SOX2 in GBM3 shows lack of responsiveness to BMP4 after 3 days of treatment (*i*.*e*. SOX2 levels are unaffected). No GFs = growth factors withdrawal. B-C) Immunostaining (B) and quantification (C) of pSMAD1,5,8 in response to BMP4 treatment in 104A (BMP-responsive) and GBM3 (BMP-resistant) GPC lines. D) qPCR quantification of the reported BMP receptors in BMP responsive (104A,PBT24,GBM2) and unresponsive (GBM3) lines. Note that GBM3 lacks expression of BMPR1A. E-F) Immunostaining (E) and quantification (F) of SOX2 staining in GBM3 GPCs electroporated with GFP control or SOX2-IRES-GFP vectors and cultured in the presence or the absence of BMP4 for 48 hours. Note that re-introducing BMPR1A in GBM3 re-establish sensitivity to BMP treatment in this line. Arrows point to GFP-expressing electroporated cells. G) qPCR quantification of SMAD4 mRNA in GBM2 cells transfected with control (siCTRL) or SMAD4 (siSMAD4) siRNAs, 3 days after transfection. H-I) Immunostaining (H) and quantification (I) of SOX2 levels in control (siCTRL) and SMAD4 (siSMAD4) GBM2 cells, cultured for 24h in the presence or the absence of BMP4. Note that in such BMP-responsive line, knock-down of SMAD4 abolishes BMP-mediated SOX2 degradation. Error bars = standard error; **p<0.005. Scale bars = 50*µ*m

The canonical pathway downstream of BMPs results in activation through phosphorylation of SMAD1,5,8 which in turn bind to the co-activator SMAD4 and activate specific transcriptional networks [23]. To understand if canonical SMAD signaling is required to mediate SOX2 degradation in GBM GPCs, we knocked-down the co-activator SMAD4 in BMP responsive GPCs. Downregulation of SMAD4 (Fig. 3G) impaired the ability of BMP4 to downregulate SOX2 (Fig. 3H-I) suggesting that a canonical SMAD signaling is required for such effect.

### BMP2/4 induces SOX2 protein degradation through the proteasome system

In order to gain mechanistic insights into the process of BMP-mediated downregulation of SOX2, we analyzed the kinetics of SOX2 protein and mRNA levels during BMP4 treatment. By western blot, SOX2 protein level downregulation was evident as early as 6 hours of treatment in concomitance with an increase in GFAP expression (Fig. 4A). In stark contrast, qPCR analysis did not reveal any downregulation of SOX2 mRNA at such early time point while still revealing GFAP mRNA increase (Fig. 4B). The decrease in SOX2 mRNA became evident only at later time points (data not shown). These data suggest that an early event of BMP signaling in GPCs is represented by SOX2 protein degradation without downregulation of the transcript. The major pathways responsible for protein degradation are the ubiquitin proteasome (UPS) and the autophagy-lysosome systems. Contrary to autophagy which functions mainly in the cytoplasm, UPS is ubiquitous and can mediates the degradation of nuclear factors [24, 25]. Considering the above, we tested whether the proteasome system is responsible for BMP-mediated SOX2 degradation. We treated GBM GPCs with BMPs and blocked the UPS with the proteasome inhibitors MG132 (Fig. 4C-E) and Bortezomib (Fig. S5). As assessed by Western blot (Fig. 4C) and immunofluorescence (Fig. 4D, Fig. S5), pharmacological inhibition of the proteasome with both compounds was able to block BMP mediated SOX2 degradation without affecting BMP signaling (i.e. SMAD1,5,8 phosphorylation, Fig. 4C). These results suggest that BMP signaling requires a functional UPS system downstream of SMAD1,5,8 in order to actively degrade SOX2 protein. Taken together, these observations suggest that BMP-mediated degradation of SOX2 requires a functional canonical BMPR1A-SMAD activation pathway which ultimately results in proteasomal degradation of SOX2.

### SOX2 in GBM-GPCs controls critical known oncogenes

Despites SOX2 being considered a critical oncogene in glioblastoma, the molecular mechanisms underlying its functions in this context are poorly understood. We knocked down SOX2 and profiled the expression of known oncogenes with established critical roles in glioblastoma such as OLIG2 [26] and RAS [27] genes. We focused on such oncogenes because, despites being critical regulators of GPCs biology, similar to SOX2, these oncogenes display low rates of somatic mutation and therefore their pathological levels could be instead achieved through regulation of transcription (Fig. S7A). We observed that knock-down of SOX2 reduced the levels of OLIG2, KRAS, NRAS, and HRAS in all the tested GPC lines while MELK, a marker of GPCs, was downregulated in 3 out of 4 lines (Fig. 5A). We next correlated the levels of SOX2 downregulation achieved in different lines with the observed reduced levels of specific oncogenes. This analysis revealed that NRAS was the oncogene with the most linear correlation with SOX2 (Pearson correlation coefficient = 0.9206, Fig. 5B). To begin testing whether SOX2 directly regulates NRAS expression, we conduct chromatin immunoprecipitation-qPCR experiments. We observed a robust binding of SOX2 at genomic elements upstream of NRAS (Fig. 5C). Notably, an analysis of 542 Glioblastoma from the TCGA database revealed that SOX2 and NRAS were significantly overexpressed in cancer versus normal brain tissue, suggesting that this oncogenic pathway has a broad relevance to glioblastoma biology (Fig. S7B). Remarkably, enforced expression of exogenous SOX2 maintains high levels of NRAS and OLIG2 proteins even when GPCs were cultured in the presence of BMP4 (Fig. 5D-E) suggesting a direct and dominant effect of SOX2 on NRAS and OLIG2 expression.

**Figure 5:**
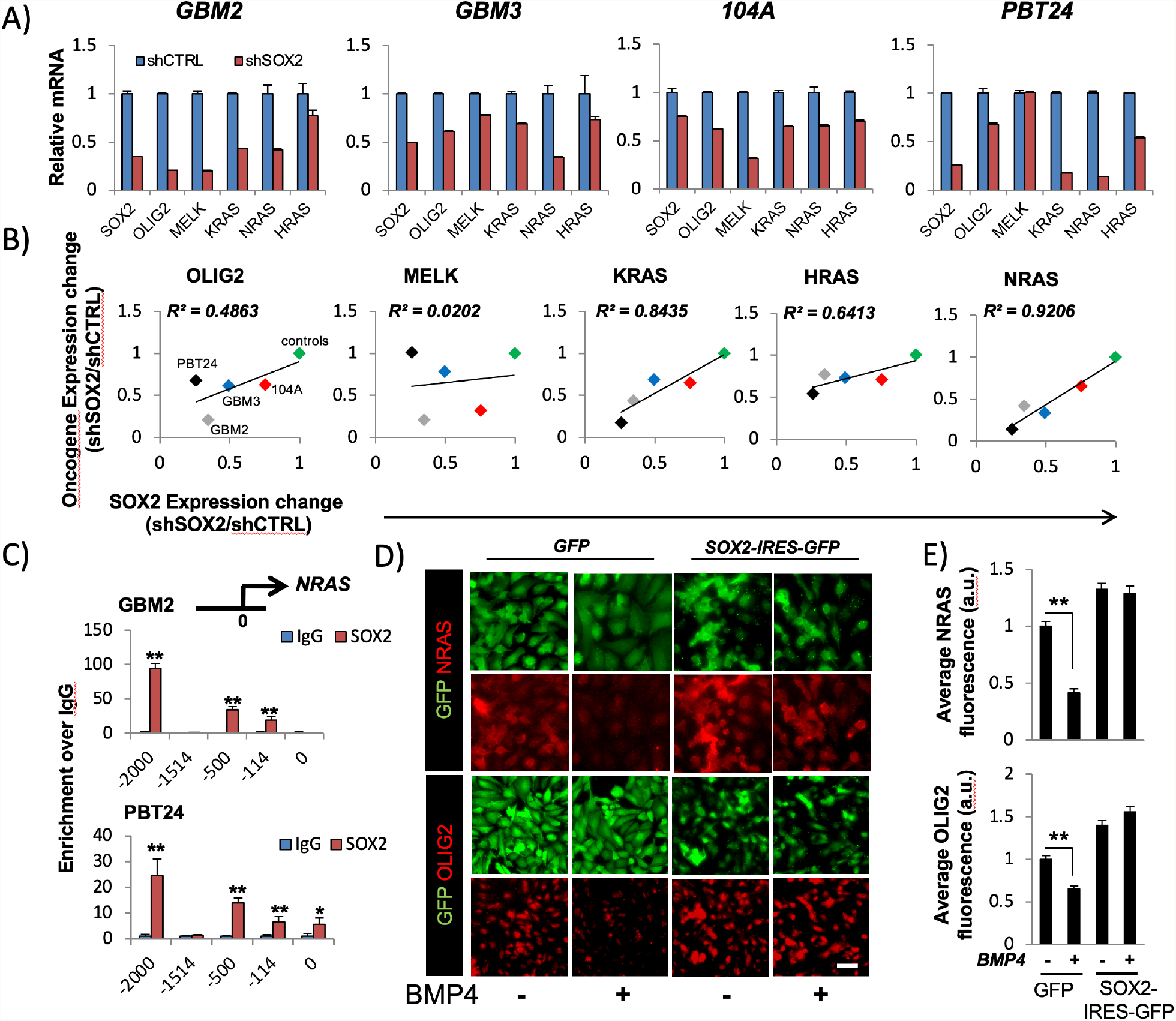
SOX2 controls oncogenes with ascertain critical role in Glioblastoma. A) qPCR for SOX2,OLIG2,MELK and RAS oncogenes in the reported GBM GPCs lines transduced with control (shCTRL) or SOX2 (shSOX2) inducible lentiviruses. SOX2 was downregulated for 4 days. B) Across lines correlation between the levels of SOX2 knock-down achieved in a particular GPC line and the resulting decrease in the reported oncogenes. Linear correlation coefficient (R^2^) is also shown. Note that NRAS displays the strongest linear dependency on SOX2 levels. Scale bars = 50*µ*m. C) Chip-qPCR showing SOX2 binding at genomic elements upstream of NRAS gene in GBM2 and PBT24 GPC lines. Numbers are relative to the transcription starting site. D) Immunostaining for NRAS and OLIG2 in PBT24 GPCs transduced with either pgk-GFP or pgk-SOX2-IRES-GFP lentiviruses and cultured in the presence or the absence of BMP4. for 3 days. E) Quantification of NRAS and OLIG2 fluorescence in the conditions reported in (D). Error bars = standard error; **p<0.005.

Overall, our results suggest that the BMP-SOX2 axis regulates a network of important players in glioblastoma biology and that manipulation of SOX2 levels can be exploited to mitigate multiple oncogenic signals in a wide range of Glioblastoma.

## DISCUSSION

A subpopulation of cancer cells with stem-like characteristics have been identified so far in several cancers such as breast, colon, lung and brain [12, 16, 28-30]. These cells have high tumorigenic capacity, resemble tissue-specific stem cells and seem to be resistant to standard chemo- and radiotherapy [14, 15]. Contrary to the idea of quiescent glioblastoma stem cells that resist therapy [4], single cell analysis of freshly dissociated tumors revealed glioblastoma propagating cell are highly proliferative [16]. Critically, RNA velocity analysis revealed conservative differentiation trajectories from GPCs to neuronal, oligo, astrocytic, and mesenchymal cells within all primary tumors analyzed [16]. Such universally present differentiation observed in freshly isolated human glioblastoma samples strongly support the idea of further promoting naturally occurring differentiating of glioblastoma GPCs into non-tumorigenic cells.

Supporting the similarities between normal neural stem cells and GPCs, we observed that genes involved in regulation of neural stem cells during development also play a critical role in GPCs. Critically, SOX2, a well-known pluripotency gene, one of 4 Yamanaka factors [31], and key regulator of NSC and embryonic stem cells biology in general [6, 7, 17, 18, 32], has been shown function as an oncogene in several GPCs from different types of tumors [20, 33-35]. BMPs are important morphogens/developmental regulators with a broad effect on normal NSCs which seem to be dependent on the developmental stage and environmental milieu [22, 36]. Here we have documented that BMP2/4 antagonize SOX2 in GPCs by inducing a rapid degradation of SOX2 protein. This effect resemble BMP4 morphogenic activity on normal neuroepithelial cells of the dorsal part of the neural tube where, in order to acquire a migratory neural crest fate, NSCs must downregulate SOX2 [6]. The ability of BMP to downregulate on SOX2 in GPCs requires both a functional proteasome system and a canonical SMAD signaling. Of note, proteasome inhibition has been proposed as a possible strategy to cancer and gliomas in particular [37], but their advance is hampered by generalized toxicity issues, likely due to a universal requirement for proteasomal degradation for organismal homeostasis. In the case of GPCs, blocking proteasome activity could prevent SOX2 degradation and differentiation of GPCs within the tumor mass. This observation might explain, at least in part, why several proteasome inhibitors failed to show efficacy in clinical trials in glioma patients [38, 39]. In mouse ES cells precise level of Sox2 protein is control by a balanced methylation and phosphorylation switch [40]. Namely, Set7 monomethyl at K119 inhibits Sox2 transcriptional activity and induces Sox2 ubiquitination and degradation. The E3 ligase WWP2 specifically interacts with K119-monomethylated Sox2 through its HECT domain to promote Sox2 ubiquitination and degradation. In contrast, AKT1 phosphorylates Sox2 at T118 and stabilizes Sox2 by antagonizing K119me by Set7 and vice versa [40]. Interestingly, competitive binding of E3 ligases TRIM26 and WWP2 was reporter to control ubiquitin proteasomal degradation of SOX2 protein in glioblastoma [41]. It is likely that BMP2/4 signaling tip the balance towards increased degradation of SOX2 through WWP2-dependent mechanism.

One of our lines (GBM3) lacked expression of BMPR1A and re-introduction of this particular BMP receptor was sufficient to re-establish sensitivity to BMP2/4 induced degradation of SOX2. Curiously, a BMP2/4-resistant glioblastoma line lacking BMPR1B expression but expressing BMPR1A was previously reported by Lee and colleagues [1]. This suggest that different receptors may mediate BMP activity in different glioblastoma lines and that different patient-specific genetic alterations might explain resistance to BMP2/4.

Despites SOX2 being known to play a critical role in maintaining tumorigenicity of glioblastoma, the molecular mechanisms underlying this effect still poorly understaood. Here we linked levels of SOX2 to the expression of several other oncogenes that share with SOX2 two main characteristics: 1) low rate of mutations in Glioblastoma and 2) a reported critical role in regulation of glioblastoma tumorigenicity. Such oncogenes (OLIG2 and RAS family genes) robustly responded to SOX2 knock-down across different GPC lines by reducing levels of expression. In particular, the connection between SOX2 and NRAS seems clinically relevant. In fact, analysis of tumors from The Cancer Genome Atlas showed robust overexpression of both oncogenic proteins across 541 glioblastoma, thus suggesting that these two oncogenes might have a general relevance in glioblastoma despites the known heterogeneity of this tumor [42, 43]. Concluding, here we identified a BMP-SOX2 axis in glioblastoma multiforme that pave the road to the development of potentially useful therapeutic treatments for the modulation of SOX2 oncogenic signaling in glioblastoma. Future experiments will focus on genes and network interaction downstream of SOX2 and NRAS as well as investigating whether proteasomal degradation of SOX2 is specific to BMP2/4 or common multiple glycogenic signals.

## EXPERIMENTAL PROCEDURES

### Cell Culture

All primary glioblastoma cell liens were established in the laboratory of Dr. Kornblum at UCLA under IRB#11-000432 Detection of Neural Stem Cells in Adult Brain Tumors.

Monolayer cultures of patient-derived GPCs cells were propagated onto Matrigel-coated plates (BD Biosciences, final concentration = 1:30, overnight coating at 4°C) using base medium (1:1 ratio of DMEM/F12 Glutamax/Neurobasal medium [Gibco], 2% B27 supplement without vitamin A [Gibco], 10% BIT 9500 [StemCell Technologies], and 1 mM glutamine [Gibco]) supplemented with 20ng/ml EGF (Chemicon), 20ng/ml bFGF, 5 µg/ml insulin (Sigma) and 5 mM nicotinamide (Sigma)). glioblastoma-GPCs were split enzymatically with Accutase (Chemicon) approximately once a week at a 1:3 ratio. For BMP/TGFβ1 treatments (all from R&D), BMP2 was used at 200ng/ml (corresponding to 1X its Effective Dosage 50 -ED50, as reported by the manufacturer), BMP4 was used at 100ng/ml (3.3X its ED50) and TGFβ1 was used up to 20 ng/ml (100X its ED50).

### Immunocytochemistry

Cells were rinsed with PBS and fixed in 4% paraformaldehyde (PFA)/PBS for 10 min at room temperature. After blocking with PBSAT (3% BSA and 0.5% Triton X-100 diluted in PBS) for 1h at room temperature, cells were incubated overnight at 4°C with the primary antibodies diluted in PBSAT. Antibodies: mouse anti-SOX2 (R&D, MAB2018), rabbit anti-GFAP (Millipore AB5804), rabbit anti-KI67 (Vector, VP-RM04), rabbit anti-OLIG2 (Millipore AB9610), mouse anti-NRAS (Santa Cruz, sc-31), pSMAD1,5,8 (Cell Signaling, 9511) The appropriate fluorochrome-conjugated secondary antibody was used with every primary antibody at a 1:1000 dilution. Nuclear co-staining was performed by incubating cells with DAPI nuclear dye.

### Quantitative PCR

Total RNA was extracted using the RNeasy kit (Qiagen) and 1 µg total RNA was reverse transcribed using the Quantitect kit (Qiagen), according to the manufacturer’s instructions. 2 µl of purified cDNA diluted 1:10 was used as a template. qPCR was performed with SYBRGreen master mix (Invitrogen) according to the manufacturer’s recommendations. HPRT or GAPDH were used for normalization. Data were analyzed using the Δ(ΔCT) method. Primers used are listed in Table S1.

### ChIP and qPCR

ChIP was performed using the Ez-ChIP kit (Millipore) according to the manufacturer’s recommendations with the following modifications: 1×10^6^ cells equivalents were used for each immunoprecipitation, cells were sonicated to yield chromatin fragments of 100-1000 bp, 5-10 µg of immunoprecipitating antibodies were employed in each ChIP. Antibodies were: normal rabbit IgG (Millipore, PP64; used a non-specific control) and rabbit anti-SOX2 (Millipore, AB5603). To evaluate factor-specific enrichment at different promoter sites, qPCR was performed using the purified chromatin as a template. Amplification was performed with site specific primers designed to flank the genomic region of interest. For a primer list see Table S1. qPCR data were normalized to the values obtained with the non-specific antibody (normal rabbit IgG).

### Lentivirus-mediated shRNA and overexpression

For shRNA experiments, lentivectors expressing doxycyclin-inducible human SOX2 shRNA or scrambled (SCR) shRNA were described before [6]. GPCs were infected with lentiviral particles to obtain lines stably carrying either SOX2 or SCR inducible shRNA. Alternatively, for short terms experiments, GPCs were acutely infected with the lentiviruses and shRNA was induced no earlier than 3 days post-infection. For SOX2 overexpression, human SOX2 coding sequence was amplified from cDNA obtained from hES-derived neural stem cells and cloned into a lentivector under control of the PGK promoter. SOX2 CDS was inserted upstream of a IRES-GFP sequence. Primers, vectors and details about the cloning strategy are available upon request. All lentivectors were packaged into lentiviral particles at the Viral Vectors Facility at the Sanford-Burnham Medical Research Institute (La Jolla, CA).

### Electroporation

BMPR1A coding sequence was inserted downstream of PGK promoter into a PGK-IRES-GFP vector. A PGK-GFP plasmid was used as a control. The vectors were electroporated into GBM3 with the Neon transfection system (Life Technologies) according to the manufacturer instructions. In details, 1·10^5^ cells were electroporated with 1.5 µg of plasmid using 3 pulses of 10ms at 1400V.

### Knock-down studies with siRNAs

Control and pooled SMAD4 siRNAs were from Dharmacon. siRNAs Were delivered with the RNAiMAX kit according to the manufacturer instructions. In details, 6·10^4^ cells were transfected using 0.5 µl of RNAiMAX reagent and 4 µl of 0.5µM siRNAs.

### Digital image analysis

For each given marker, pictures of cells cultured under different experimental conditions were taken with the same exposure time and contrast/brightness parameters. When amount of fluorescence was evaluated, the software ImageJ was used (http://imagej.nih.gov/ij/). A minimum of 3 random fields containing at least 100 cells was analyzed for each condition.

## Acknowledgements

We thank Sanford Burnham Prebys microscopy and pathology Core Facilities for the help with sample preparations and image acquisition.

---

**Figure S1:**
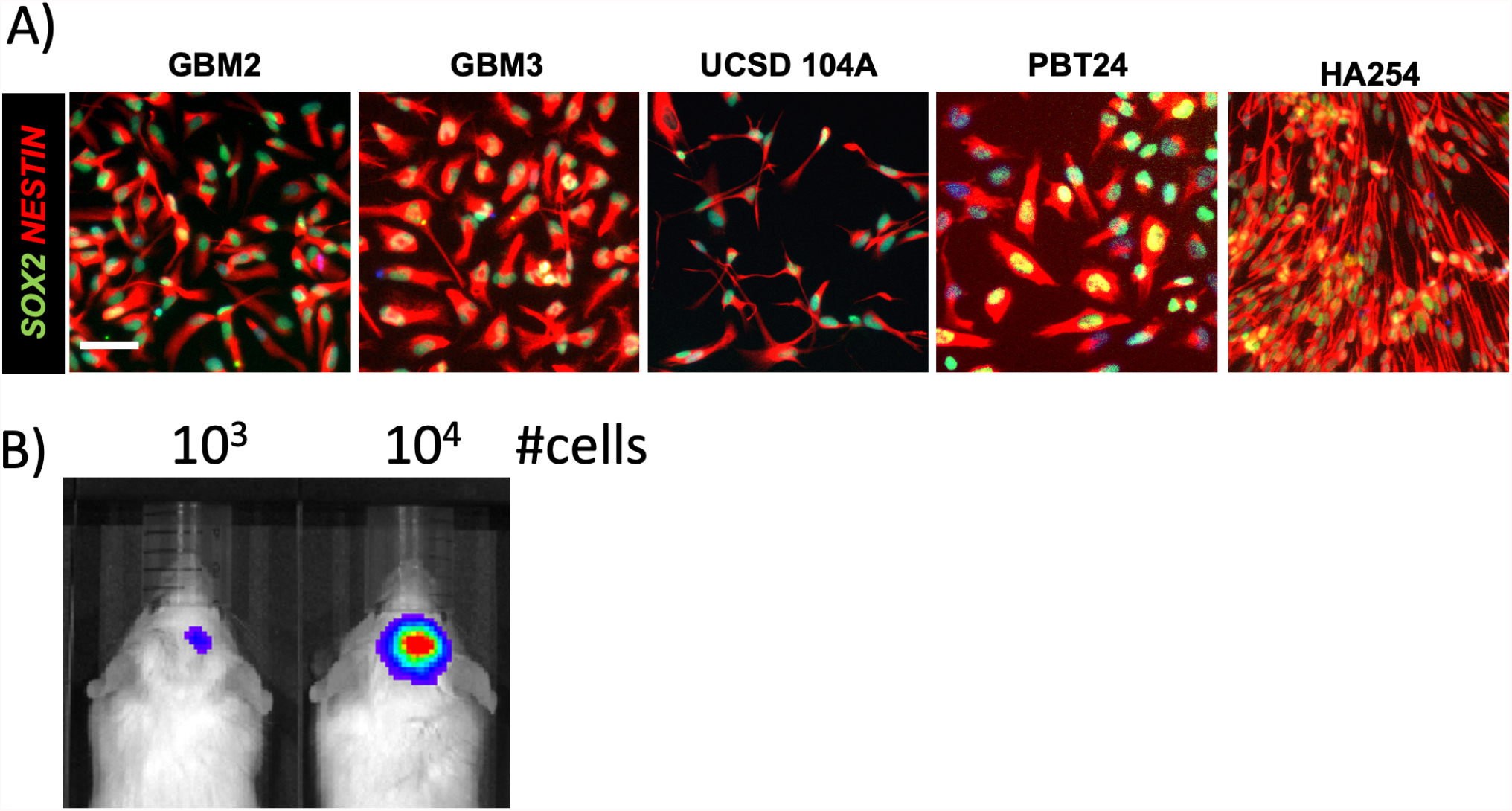
Human patient-derived GLIOBLASTOMA GPCs. A) immunostaining for the neural stem cell markers SOX2 and NESTIN in 5 patient-derived glioblastoma GPCs. Scale bar = 50µm B) Xenografts of PBT24 GPCs transduced with luciferase lentivirus showing tumors 42 days after transplantation in NSG immunodeficient mice. Number of transplanted cells is reported in the pictures. Cells were stereo tactically injected in the right hippocampus.

**Figure S2:**
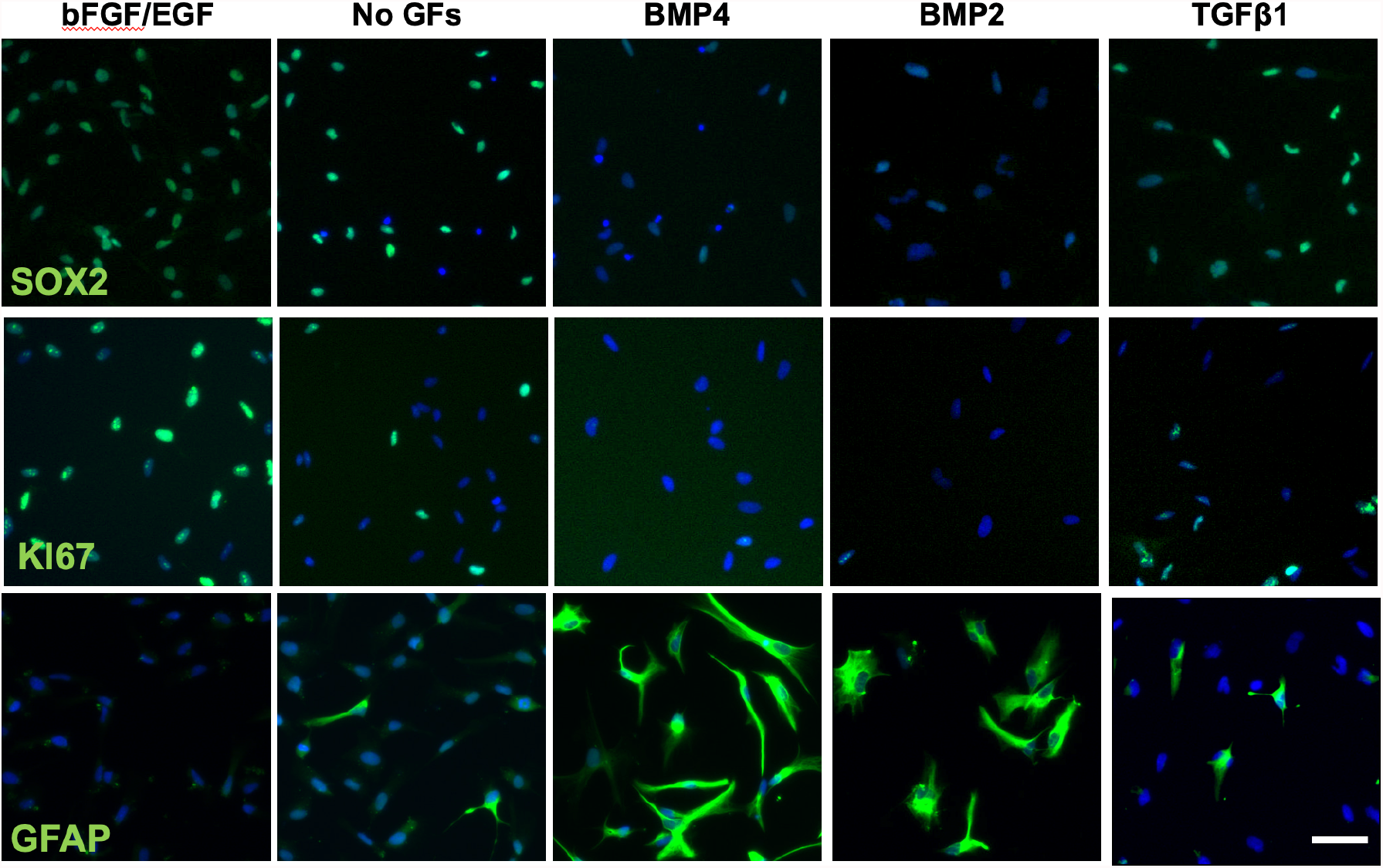
Effect of different treatments on glioblastoma GPCs. A) immunostaining for SOX2, KI67 and GFAP in GBM2 GPCs cultured for 3 days with bFGF/EGF mitogens, growth factor withdrawal (No GFs), BMP2, BMP4 or TGFβ1. Scale bar = 50*µ*m

**Figure S3:**
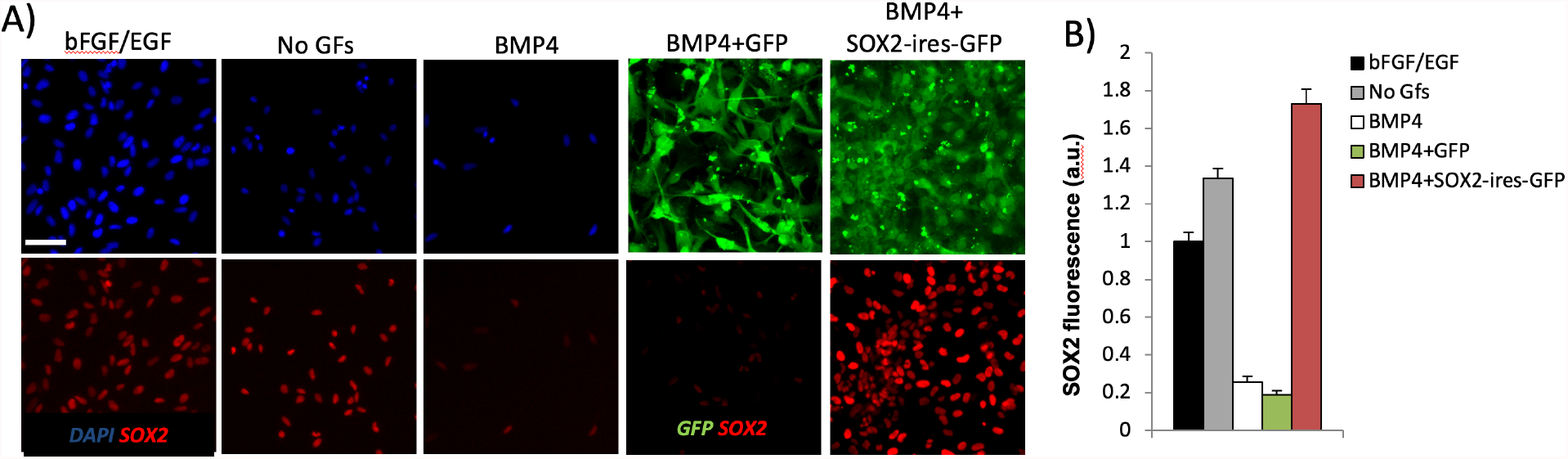
Stable SOX2 overexpression in BMP treated GPCs. A) immunostaining for SOX2 in BMP4 treated GBM2 GPCs (3 days of treatment) infected with pgk-GFP or pgk-SOX2-IRES-GFP lentiviruses. Wild type cells treated with bFGF/EGF, no growth factors (NoGFs) or BMP4 are included for comparison. Scale bar = 50*µ*m. B) Quantification of SOX2 levels for conditions shown in (A).

**Figure S4:**
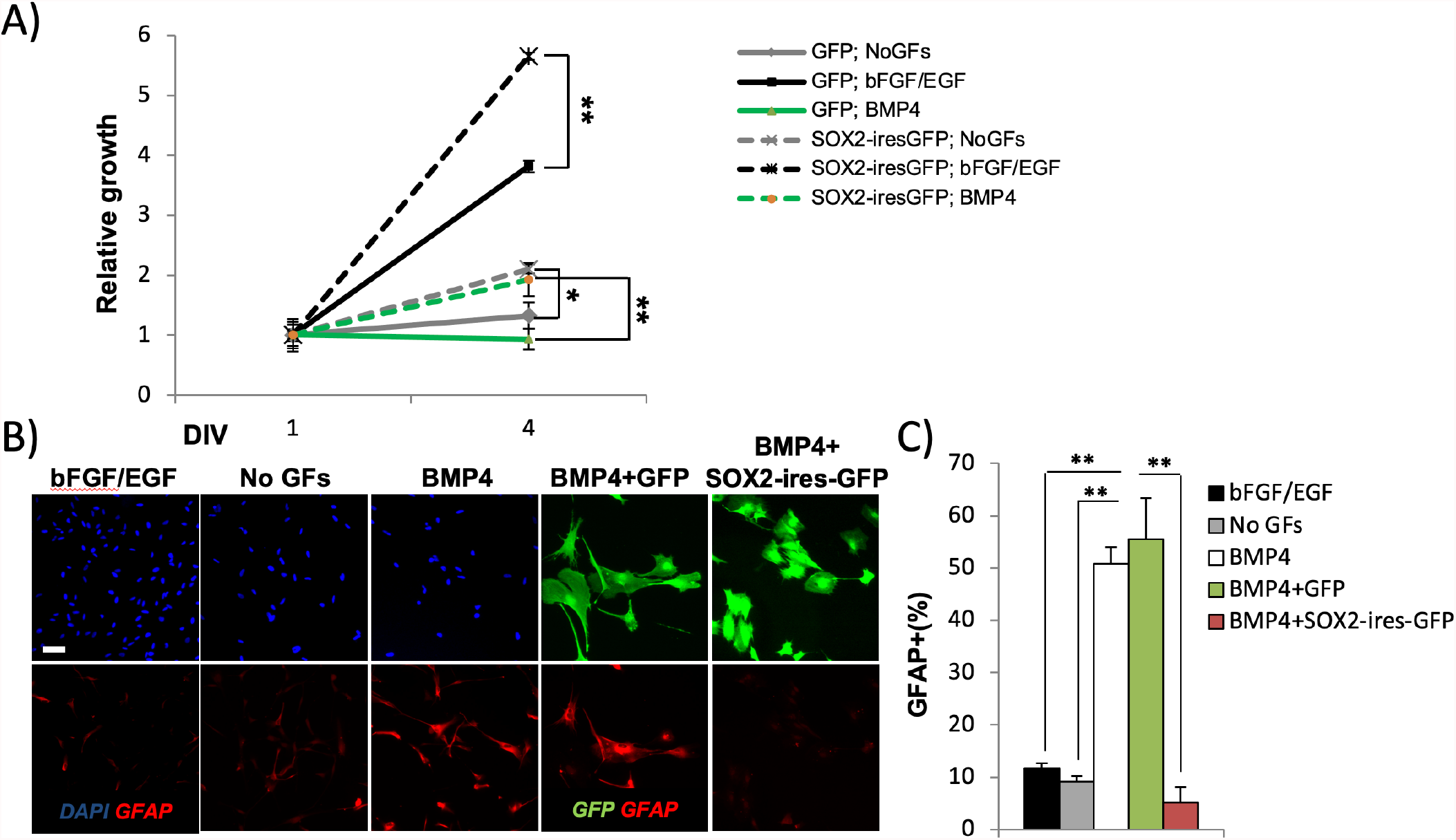
SOX2 rescues BMP effect on GPCs. A) Growth curve of 104A GPCs infected with pgk-GFP or pgk-SOX2-IRES-GFP and cultured in the presence of bFGF/EGF mitogens, no growth factors (No GFs) or BMP4 for 4 days. Note that SOX2 overexpression overcomes the oncostatic effect of BMP4 (comparison of dashed and solid green curves). B-C) Immunostaining (B) and quantification (C) of GFAP in BMP2/4 treated 104A GPCs (3 days of treatment) infected with pgk-GFP or pgk-SOX2-IRES-GFP lentiviruses. Note that SOX2 overexpression abolishes BMP4-mediated increase in gliogenesis. Error bars = standard error; *p<0.05; **p<0.005. Scale bar = 50*µ*m

**Figure S5:**
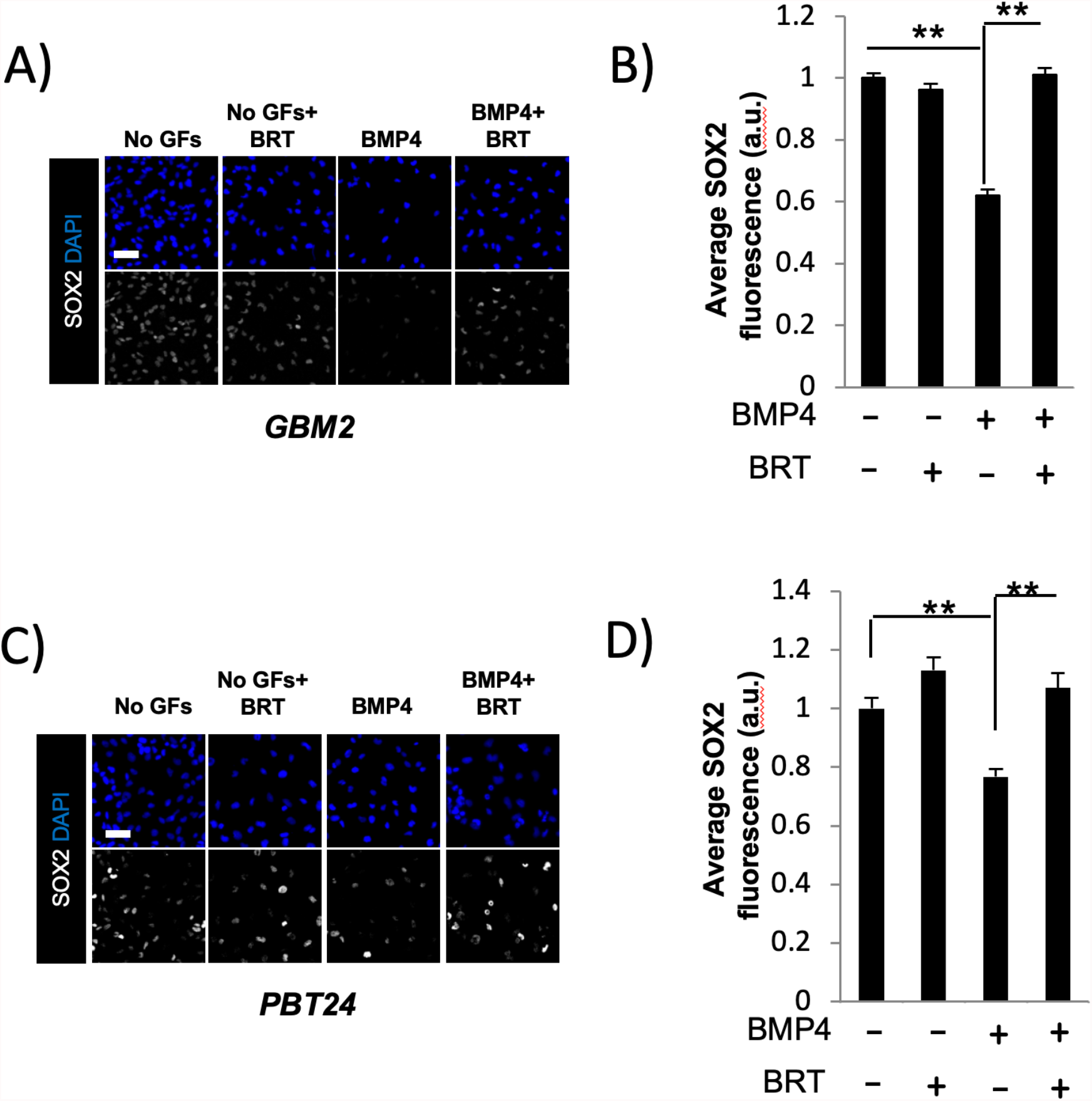
The proteasome inhibitor Bortezomib blocks BMP-mediated SOX2 degradation. A-B) Immunostaining (A) and quantification (B) of SOX2 in GBM2 GPCs cultured under the reported conditions for 18 hours. Note that bortezomib (BRT) blocks degradation of SOX2 in the presence of BMP4. C-D) Immunostaining (C) and quantification (D) of SOX2 in PBT24 GPCs cultured under the reported conditions for 18 hours. Error bars = standard error; **p<0.005. Scale bar = 50*µ*m

**Figure S6:**
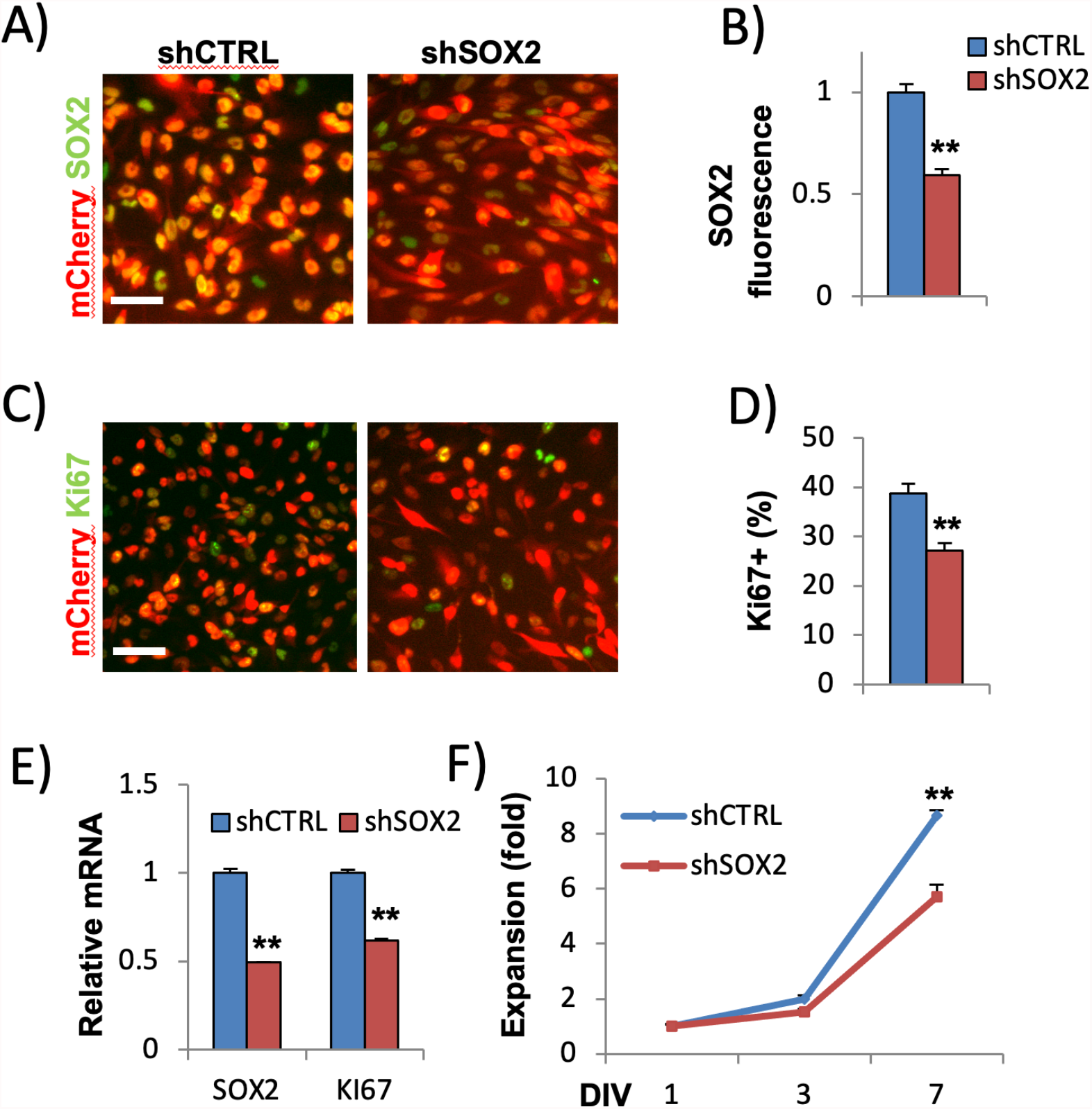
BMP-resistant GBM3 is sensitive to SOX2 knockdown. A-B) Immunostaining (A) and quantification (B) of SOX2 in GBM3 transduced with doxycycline-inducible control (shCTRL) and SOX2 (shSOX2) shRNA. mCherry labels transduced cells. shRNA was induced for 10 days. C-D) Immunostaining (A) and quantification (B) of KI67 in GBM3 transduced with doxycycline-inducible control (shCTRL) and SOX2 (shSOX2) shRNA. (E) qPCR confirming SOX2 and KI67 downregulation in shSOX2 versus shCTRL GBM3. F) Growth curve showing reduced proliferation of shSOX2 versus shCTRL GBM3. Error bars = standard error; **p<0.005. Scale bars = 50*µ*m

**Figure S7:**
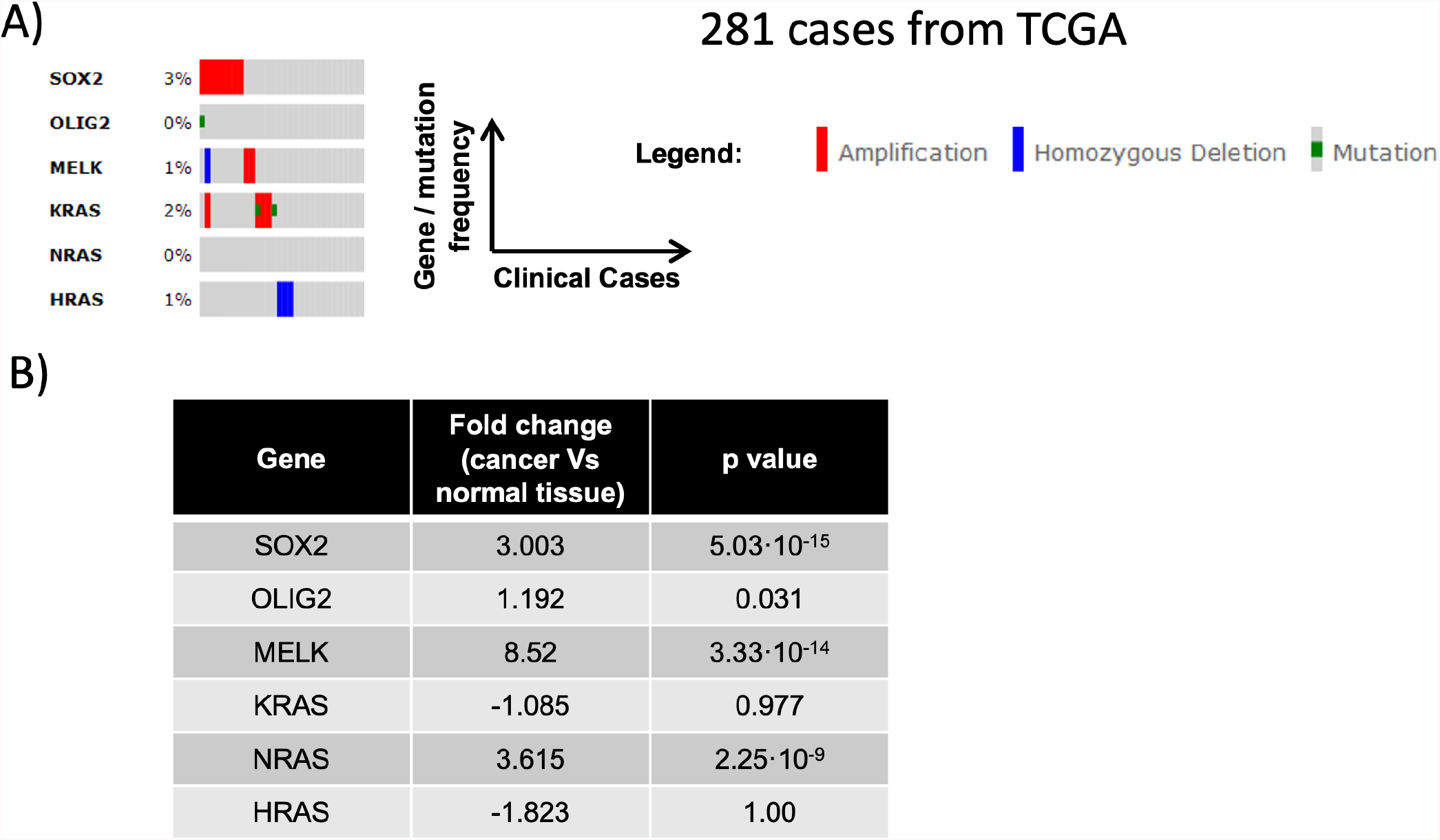
Bioinformatics analysis of TCGA glioblastoma tumors. A) Analysis of 281 TCGA glioblastoma clinical cases with known copy number alterations and mutations. Each column represents an individual case. Percentage and type of mutations for SOX2, OLIG2, MELK and RAS genes are reported. Analysis was performed *via* the cBIO portal (http://www.cbioportal.org/public-portal/) B) Analysis of expression of SOX2, OLIG2, MELK and RAS oncogenes in 541 TCGA glioblastoma lesions versus normal brain tissue. Analysis was performed *via* the the Oncomine portal (https://www.oncomine.org/resource/login.html).

**Table S1.**
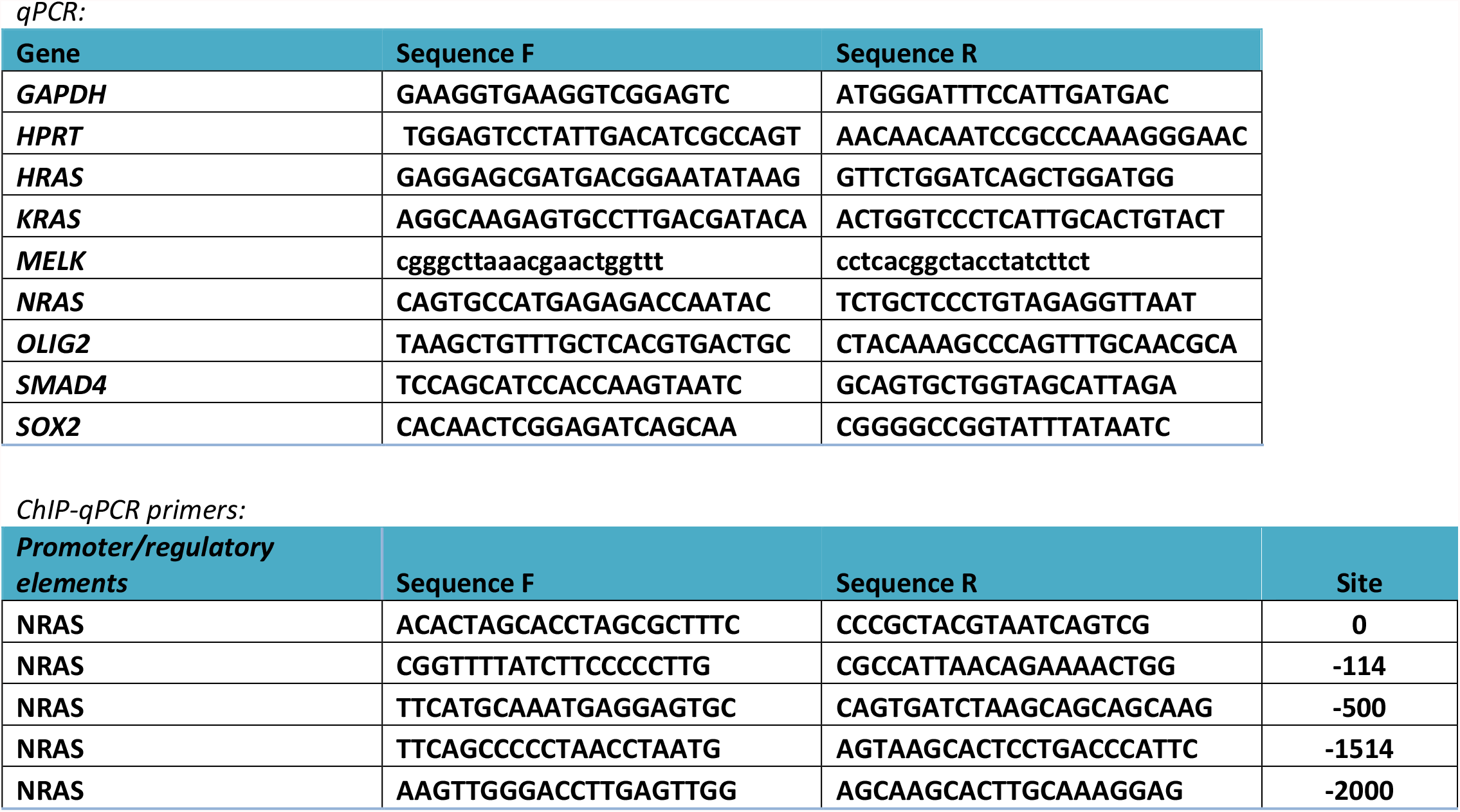

## References

1. Lee J, Son MJ, Woolard K, Donin NM, Li A, Cheng CH, Kotliarova S, Kotliarov Y, Walling J, Ahn S, Kim M, Totonchy M, Cusack T, Ene C, Ma H, Su Q, Zenklusen JC, Zhang W, Maric D, Fine HA. Epigenetic-mediated dysfunction of the bone morphogenetic protein pathway inhibits differentiation of glioblastoma-initiating cells. Cancer Cell. 2008;13(1):69–80. Epub 2008/01/03. doi: 10.1016/j.ccr.2007.12.005. PubMed PMID: 18167341; PMCID: 2835498.

2. Piccirillo SG, Reynolds BA, Zanetti N, Lamorte G, Binda E, Broggi G, Brem H, Olivi A, Dimeco F, Vescovi AL. Bone morphogenetic proteins inhibit the tumorigenic potential of human brain tumour-initiating cells. Nature. 2006;444(7120):761–5. Epub 2006/12/08. doi: 10.1038/nature05349. PubMed PMID: 17151667.

3. Tate CM, Pallini R, Ricci-Vitiani L, Dowless M, Shiyanova T, D’Alessandris GQ, Morgante L, Giannetti S, Larocca LM, di Martino S, Rowlinson SW, De Maria R, Stancato L. A BMP7 variant inhibits the tumorigenic potential of glioblastoma stem-like cells. Cell Death Differ. 2012;19(10):1644–54. PubMed PMID: 22539003.

4. Sachdeva R, Wu M, Johnson K, Kim H, Celebre A, Shahzad U, Graham MS, Kessler JA, Chuang JH, Karamchandani J, Bredel M, Verhaak R, Das S. BMP signaling mediates glioma stem cell quiescence and confers treatment resistance in glioblastoma. Sci Rep. 2019;9(1):14569. Epub 2019/10/12. doi: 10.1038/s41598-019-51270-1. PubMed PMID: 31602000; PMCID: PMC6787003.

5. Dalmo E, Johansson P, Niklasson M, Gustavsson I, Nelander S, Westermark B. Growth-Inhibitory Activity of Bone Morphogenetic Protein 4 in Human Glioblastoma Cell Lines Is Heterogeneous and Dependent on Reduced SOX2 Expression. Mol Cancer Res. 2020;18(7):981–91. Epub 2020/04/03. doi: 10.1158/1541-7786.MCR-19-0638. PubMed PMID: 32234828.

6. Cimadamore F, Fishwick K, Giusto E, Gnedeva K, Cattarossi G, Miller A, Pluchino S, Brill LM, Bronner-Fraser M, Terskikh AV. Human ESC-Derived Neural Crest Model Reveals a Key Role for SOX2 in Sensory Neurogenesis. Cell Stem Cell. 2011;8(5):538–51. Epub 2011/05/10. doi: S1934-5909(11)00121-4 [pii]10.1016/j.stem.2011.03.011. PubMed PMID: 21549328.

7. Cimadamore F, Amador-Arjona A, Chen C, Huang CT, Terskikh AV. SOX2-LIN28/let-7 pathway regulates proliferation and neurogenesis in neural precursors. Proceedings of the National Academy of Sciences of the United States of America. 2013;110(32):E3017–26. Epub 2013/07/26. doi: 10.1073/pnas.1220176110. PubMed PMID: 23884650; PMCID: 3740872.

8. Amador-Arjona A, Cimadamore F, Huang CT, Wright R, Lewis S, Gage FH, Terskikh AV. SOX2 primes the epigenetic landscape in neural precursors enabling proper gene activation during hippocampal neurogenesis. Proc Natl Acad Sci U S A. 2015;112(15):E1936–45. Epub 2015/04/01. doi: 10.1073/pnas.1421480112. PubMed PMID: 25825708; PMCID: 4403144.

9. Porter KR, McCarthy BJ, Freels S, Kim Y, Davis FG. Prevalence estimates for primary brain tumors in the United States by age, gender, behavior, and histology. Neuro Oncol. 2010;12(6):520–7. PubMed PMID: 20511189.

10. Stupp R, Mason WP, van den Bent MJ, Weller M, Fisher B, Taphoorn MJ, Belanger K, Brandes AA, Marosi C, Bogdahn U, Curschmann J, Janzer RC, Ludwin SK, Gorlia T, Allgeier A, Lacombe D, Cairncross JG, Eisenhauer E, Mirimanoff RO. Radiotherapy plus concomitant and adjuvant temozolomide for glioblastoma. N Engl J Med. 2005;352(10):987–96. PubMed PMID: 15758009.

11. Lee J, Kotliarova S, Kotliarov Y, Li A, Su Q, Donin NM, Pastorino S, Purow BW, Christopher N, Zhang W, Park JK, Fine HA. Tumor stem cells derived from glioblastomas cultured in bFGF and EGF more closely mirror the phenotype and genotype of primary tumors than do serum-cultured cell lines. Cancer Cell. 2006;9(5):391–403. Epub 2006/05/16. doi: 10.1016/j.ccr.2006.03.030. PubMed PMID: 16697959.

12. Singh SK, Hawkins C, Clarke ID, Squire JA, Bayani J, Hide T, Henkelman RM, Cusimano MD, Dirks PB. Identification of human brain tumour initiating cells. Nature. 2004;432(7015):396–401. PubMed PMID: 15549107.

13. Galli R, Binda E, Orfanelli U, Cipelletti B, Gritti A, De Vitis S, Fiocco R, Foroni C, Dimeco F, Vescovi A. Isolation and characterization of tumorigenic, stem-like neural precursors from human glioblastoma. Cancer Res. 2004;64(19):7011–21. PubMed PMID: 15466194.

14. Bao S, Wu Q, McLendon RE, Hao Y, Shi Q, Hjelmeland AB, Dewhirst MW, Bigner DD, Rich JN. Glioma stem cells promote radioresistance by preferential activation of the DNA damage response. Nature. 2006;444(7120):756–60. PubMed PMID: 17051156.

15. Chen J, Li Y, Yu TS, McKay RM, Burns DK, Kernie SG, Parada LF. A restricted cell population propagates glioblastoma growth after chemotherapy. Nature. 2012;488(7412):522–6. PubMed PMID: 22854781.

16. Couturier CP, Ayyadhury S, Le PU, Nadaf J, Monlong J, Riva G, Allache R, Baig S, Yan X, Bourgey M, Lee C, Wang YCD, Wee Yong V, Guiot MC, Najafabadi H, Misic B, Antel J, Bourque G, Ragoussis J, Petrecca K. Single-cell RNA-seq reveals that glioblastoma recapitulates a normal neurodevelopmental hierarchy. Nat Commun. 2020;11(1):3406. Epub 2020/07/10. doi: 10.1038/s41467-020-17186-5. PubMed PMID: 32641768; PMCID: PMC7343844.

17. Cavallaro M, Mariani J, Lancini C, Latorre E, Caccia R, Gullo F, Valotta M, DeBiasi S, Spinardi L, Ronchi A, Wanke E, Brunelli S, Favaro R, Ottolenghi S, Nicolis SK. Impaired generation of mature neurons by neural stem cells from hypomorphic Sox2 mutants. Development. 2008;135(3):541–57. PubMed PMID: 18171687.

18. Graham V, Khudyakov J, Ellis P, Pevny L. SOX2 Functions to Maintain Neural Progenitor Identity. Neuron. 2003;39(5):749–65. PubMed PMID: 12948443.

19. Annovazzi L, Mellai M, Caldera V, Valente G, Schiffer D. SOX2 expression and amplification in gliomas and glioma cell lines. Cancer Genomics Proteomics. 2011;8(3):139–47. PubMed PMID: 21518820.

20. Gangemi RM, Griffero F, Marubbi D, Perera M, Capra MC, Malatesta P, Ravetti GL, Zona GL, Daga A, Corte G. SOX2 silencing in glioblastoma tumor-initiating cells causes stop of proliferation and loss of tumorigenicity. Stem Cells. 2009;27(1):40–8. PubMed PMID: 18948646.

21. Lim HN, Berkovitz GD, Hughes IA, Hawkins JR. Mutation analysis of subjects with 46, XX sex reversal and 46, XY gonadal dysgenesis does not support the involvement of SOX3 in testis determination. Hum Genet. 2000;107(6):650–2. Epub 2001/01/12. doi: 10.1007/s004390000428. PubMed PMID: 11153920.

22. Mehler MF, Mabie PC, Zhu G, Gokhan S, Kessler JA. Developmental changes in progenitor cell responsiveness to bone morphogenetic proteins differentially modulate progressive CNS lineage fate. Dev Neurosci. 2000;22(1-2):74–85. PubMed PMID: 10657700.

23. Miyazawa K, Shinozaki M, Hara T, Furuya T, Miyazono K. Two major Smad pathways in TGF-beta superfamily signalling. Genes Cells. 2002;7(12):1191–204. PubMed PMID: 12485160.

24. Li XJ, Li S. Proteasomal dysfunction in aging and Huntington disease. Neurobiol Dis. 2011;43(1):4–8. PubMed PMID: 21145396.

25. von Mikecz A. The nuclear ubiquitin-proteasome system. J Cell Sci. 2006;119(Pt 10):1977-84. PubMed PMID: 16687735.

26. Ligon KL, Huillard E, Mehta S, Kesari S, Liu H, Alberta JA, Bachoo RM, Kane M, Louis DN, Depinho RA, Anderson DJ, Stiles CD, Rowitch DH. Olig2-regulated lineage-restricted pathway controls replication competence in neural stem cells and malignant glioma. Neuron. 2007;53(4):503–17. PubMed PMID: 17296553.

27. Holland EC, Celestino J, Dai C, Schaefer L, Sawaya RE, Fuller GN. Combined activation of Ras and Akt in neural progenitors induces glioblastoma formation in mice. Nat Genet. 2000;25(1):55–7. PubMed PMID: 10802656.

28. Eramo A, Lotti F, Sette G, Pilozzi E, Biffoni M, Di Virgilio A, Conticello C, Ruco L, Peschle C, De Maria R. Identification and expansion of the tumorigenic lung cancer stem cell population. Cell Death Differ. 2008;15(3):504–14. PubMed PMID: 18049477.

29. O’Brien CA, Pollett A, Gallinger S, Dick JE. A human colon cancer cell capable of initiating tumour growth in immunodeficient mice. Nature. 2007;445(7123):106–10. PubMed PMID: 17122772.

30. Liu S, Dontu G, Wicha MS. Mammary stem cells, self-renewal pathways, and carcinogenesis. Breast Cancer Res. 2005;7(3):86–95. PubMed PMID: 15987436.

31. Takahashi K, Yamanaka S. Induction of pluripotent stem cells from mouse embryonic and adult fibroblast cultures by defined factors. Cell. 2006;126(4):663–76. PubMed PMID: 16904174.

32. Rizzino A. Sox2 and Oct-3/4: a versatile pair of master regulators that orchestrate the self-renewal and pluripotency of embryonic stem cells. Wires Syst Biol Med. 2009;1(2):228–36. doi: 10.1002/wsbm.12. PubMed PMID: WOS:000283709600009.

33. Justilien V, Walsh MP, Ali SA, Thompson EA, Murray NR, Fields AP. The PRKCI and SOX2 oncogenes are coamplified and cooperate to activate Hedgehog signaling in lung squamous cell carcinoma. Cancer Cell. 2014;25(2):139–51. Epub 2014/02/15. doi: 10.1016/j.ccr.2014.01.008. PubMed PMID: 24525231; PMCID: 3949484.

34. Rudin CM, Durinck S, Stawiski EW, Poirier JT, Modrusan Z, Shames DS, Bergbower EA, Guan Y, Shin J, Guillory J, Rivers CS, Foo CK, Bhatt D, Stinson J, Gnad F, Haverty PM, Gentleman R, Chaudhuri S, Janakiraman V, Jaiswal BS, Parikh C, Yuan W, Zhang Z, Koeppen H, Wu TD, Stern HM, Yauch RL, Huffman KE, Paskulin DD, Illei PB, Varella-Garcia M, Gazdar AF, de Sauvage FJ, Bourgon R, Minna JD, Brock MV, Seshagiri S. Comprehensive genomic analysis identifies SOX2 as a frequently amplified gene in small-cell lung cancer. Nat Genet. 2012;44(10):1111–6. PubMed PMID: 22941189.

35. Leis O, Eguiara A, Lopez-Arribillaga E, Alberdi MJ, Hernandez-Garcia S, Elorriaga K, Pandiella A, Rezola R, Martin AG. Sox2 expression in breast tumours and activation in breast cancer stem cells. Oncogene. 2012;31(11):1354–65. PubMed PMID: 21822303.

36. Lim DA, Tramontin AD, Trevejo JM, Herrera DG, Garcia-Verdugo JM, Alvarez-Buylla A. Noggin antagonizes BMP signaling to create a niche for adult neurogenesis. Neuron. 2000;28(3):713–26.

37. Park JE, Miller Z, Jun Y, Lee W, Kim KB. Next-generation proteasome inhibitors for cancer therapy. Transl Res. 2018;198:1–16. Epub 2018/04/15. doi: 10.1016/j.trsl.2018.03.002. PubMed PMID: 29654740; PMCID: PMC6151281.

38. Ahluwalia MS, Patton C, Stevens G, Tekautz T, Angelov L, Vogelbaum MA, Weil RJ, Chao S, Elson P, Suh JH, Barnett GH, Peereboom DM. Phase II trial of ritonavir/lopinavir in patients with progressive or recurrent high-grade gliomas. J Neurooncol. 2011;102(2):317–21. PubMed PMID: 20683757.

39. Friday BB, Anderson SK, Buckner J, Yu C, Giannini C, Geoffroy F, Schwerkoske J, Mazurczak M, Gross H, Pajon E, Jaeckle K, Galanis E. Phase II trial of vorinostat in combination with bortezomib in recurrent glioblastoma: a north central cancer treatment group study. Neuro Oncol. 2012;14(2):215–21. PubMed PMID: 22090453.

40. Fang L, Zhang L, Wei W, Jin XL, Wang P, Tong YF, Li JW, D. JX, Wong JM. A Methylation-Phosphorylation Switch Determines Sox2 Stability and Function in ESC Maintenance or Differentiation. Molecular Cell. 2014;55(4):537–51. doi: 10.1016/j.molcel.2014.06.018. PubMed PMID: WOS:000340612700005.

41. Mahlokozera T, Patel B, Chen H, Desouza P, Qu X, Mao DD, Hafez D, Yang W, Taiwo R, Paturu M, Salehi A, Gujar AD, Dunn GP, Mosammaparast N, Petti AA, Yano H, Kim AH. Competitive binding of E3 ligases TRIM26 and WWP2 controls SOX2 in glioblastoma. Nat Commun. 2021;12(1):6321. Epub 2021/11/05. doi: 10.1038/s41467-021-26653-6. PubMed PMID: 34732716; PMCID: PMC8566473.

42. Phillips HS, Kharbanda S, Chen R, Forrest WF, Soriano RH, Wu TD, Misra A, Nigro JM, Colman H, Soroceanu L, Williams PM, Modrusan Z, Feuerstein BG, Aldape K. Molecular subclasses of high-grade glioma predict prognosis, delineate a pattern of disease progression, and resemble stages in neurogenesis. Cancer Cell. 2006;9(3):157–73. PubMed PMID: 16530701.

43. Verhaak RG, Hoadley KA, Purdom E, Wang V, Qi Y, Wilkerson MD, Miller CR, Ding L, Golub T, Mesirov JP, Alexe G, Lawrence M, O’Kelly M, Tamayo P, Weir BA, Gabriel S, Winckler W, Gupta S, Jakkula L, Feiler HS, Hodgson JG, James CD, Sarkaria JN, Brennan C, Kahn A, Spellman PT, Wilson RK, Speed TP, Gray JW, Meyerson M, Getz G, Perou CM, Hayes DN. Integrated genomic analysis identifies clinically relevant subtypes of glioblastoma characterized by abnormalities in PDGFRA, IDH1, EGFR, and NF1. Cancer Cell. 2010;17(1):98–110. Epub 2010/02/05. doi: S1535-6108(09)00432-2 [pii] 10.1016/j.ccr.2009.12.020. PubMed PMID: 20129251; PMCID: 2818769.

